# Hsc70-4 mediates internalization of environmental dsRNA at the surface of *Drosophila* S2 cells

**DOI:** 10.1101/2023.03.11.532206

**Authors:** Sabrina Johanna Fletcher, Lorena Tomé-Poderti, Vanesa Mongelli, Lionel Frangeul, Hervé Blanc, Yann Verdier, Joelle Vinh, Maria-Carla Saleh

## Abstract

The siRNA pathway is the primary antiviral defense mechanism in invertebrates and plants. The systemic nature of this defense mechanism is one of its more fascinating characteristics and the recognition and transport of double-stranded RNA (dsRNA) of viral origin is required for the systemic activity of the siRNA pathway. Indeed, cellular internalization of dsRNA from the environment is a widespread phenomenon among insects. Here we aimed to identify cell surface proteins that bind to extracellular dsRNA and mediate its internalization. To this end, we developed a novel co-immunoprecipitation protocol that we followed with proteomics analysis. Among the hits from our screens was Hsc70-4, a constitutively expressed member of the heat shock protein family that has been implicated in clathrin-mediated endocytosis. We found that silencing Hsc70-4 impaired dsRNA internalization. Surprisingly, despite lacking a predicted transmembrane domain, Hsc70-4 localizes to the cell membrane and this localization was preserved when Hsc70-4 was expressed in mammalian cells, suggesting a conserved role at the cell surface. Furthermore, Hsc70-4 shows a previously undescribed dsRNA-specific binding capacity. Our results show that Hsc70-4 is a key element of the dsRNA internalization process and its detailed study may facilitate the development of RNA interference (RNAi)-based technologies for pest and vector borne disease control.

**Importance:** To protect plants from pathogens or pests, the technology of “Host-induced gene silencing” has emerged as a powerful alternative to chemical treatments. This is an RNAi-based technology where small RNAs made in the plant silence the genes of the pests or pathogens that attack the plant. The small RNAs are generally derived from dsRNA expressed in transgenic plants. Alternatively, dsRNA can be sprayed onto the plant surface, where it can be taken up into the plant or ingested by pests. We have identified a cell surface protein that mediates the early steps of extracellular dsRNA internalization in insect cells. This could facilitate the development of new strategies for pest management.

## Introduction

Systemic immunity is a defense mechanism that is triggered to fight pathogen infections while contributing to resistance to infection in non-infected cells or tissues(1–3). This type of immune response is critical to maintain homeostasis. One of the main characteristics of systemic immunity is the transmission of a signal from infected cells/tissues to non-infected cells/tissues to trigger a broader immune response(2, 3). During the infection process, conserved pathogen-associated motifs known as pathogen-associated molecular patterns (PAMPs) are recognized by a variety of pattern recognition receptors that initiate immune responses(4). For example, in *Drosophila melanogaster*, peptidoglycans from Gram-negative bacteria are sensed by peptidoglycan recognition proteins, resulting in activation of the IMD pathway to produce antimicrobial peptides(2). In vertebrates, intracellular viral dsRNA is sensed by receptors such as endosomal transmembrane toll-like receptors(5) and the RIG-I-like receptors, RIG-I and MDA5(6, 7). Recognition of dsRNA by these receptors induces a transcriptional program that generates an antiviral state characterized by the secretion of cytokines such as type I interferon that can act in autocrine and/or paracrine fashions to induce the transcription of interferon-stimulated genes. In insects, Vago has been described to have an interferon-like role(8, 9). However, RNA interference (RNAi) mediated by siRNAs is the primary and most well characterized intracellular antiviral response(10, 11). During their replication cycles, viruses produce dsRNA intermediates(12). Sensing and cleavage of virus-derived dsRNA into siRNAs by intracellular proteins triggers the RNAi-mediated immune response to control viral replication.

The antiviral siRNA pathway has been most extensively studied in the model insect, *D. melanogaster*(13–16). Specifically, *in vivo* experiments in adult flies revealed that virus-derived dsRNA is transmitted from infected cells to non-infected cells, where it is processed into siRNAs and confers non-infected cells with pre-exposure protection against Drosophila C virus (DCV) or Sindbis virus infection(17).

Even though intracellular RNAi processes are well understood, the release, transmission, detection, and internalization of dsRNA is still very poorly characterized. While the release of dsRNA from infected cells is thought to be a consequence of cell lysis caused by viral infection(17), how dsRNA is internalized by non-infected cells is poorly understood. Hemocytes isolated from both larvae and adult flies can internalize naked extracellular dsRNA, causing activation of the siRNA pathway(18, 19). Accordingly, *in vitro* experiments using the hemocyte-like *D. melanogaster* Schneider 2 (S2) cell line revealed that these cells can also internalize dsRNA by receptor-mediated endocytosis under certain conditions(20, 21). These findings imply the existence of a receptor(s) that binds naked extracellular dsRNA at the cell surface prior to internalization. Several proteins have already been proposed as dsRNA receptors in other organisms, such as the systemic RNAi-defective-2 protein (SID-2) in *C. elegans*(22), and the scavenger receptor class A (SR-A) and macrophage-1 antigen (Mac-1) in humans(23–26). Furthermore, the endosomal channel protein, SID-1, has been shown to mediate the transport of endocytized dsRNA into the cytoplasm in *C. elegans*(22). In mammals, a similar role has been identified for SIDT2, the mammalian ortholog of SID-1(27). In insects, scavenger receptors have also been found to mediate the early steps in the internalization of dsRNA(20, 28). Specifically, in *D. melanogaster*, two scavenger receptors, SR-CI and Eater(21), have been proposed, however, experimental data showed that these receptors are mainly involved in general phagocytosis, and not in the specific internalization of dsRNA by endocytosis(29).

Here, we aimed to identify the cell surface protein(s) responsible for binding extracellular dsRNA prior to its internalization in *D. melanogaster*. To this end, we developed an *in vitro* model using the S2 cell line. We used two different proteomic approaches to find candidate proteins. One approach aimed to identify cell surface proteins. The other approach was designed to specifically pull-down dsRNA-protein complexes formed at the cell surface. Through a functional screen of candidate proteins, we identified a protein, Hsc70-4, that localizes to the cell surface, binds dsRNA, and plays a previously uncharacterized role in dsRNA internalization in S2 cells.

## Results

### S2 cells internalize dsRNA when grown in a protein-free medium

*D. melanogaster* S2 cells are widely used as a tool to study gene function because of their capacity to internalize dsRNA from the cell medium to induce gene silencing through RNAi. dsRNA internalization in S2 cells depends on the composition of the growth medium and on the specific cell line used. While S2 cells internalize dsRNA from the growth medium only in the absence of fetal bovine serum (FBS), the S2 receptor plus (S2R+)(30) cells can internalize dsRNA in the presence or absence of FBS(31). With the objective of establishing an experimental model to identify the proteins involved in the internalization of dsRNA from the extracellular milieu, we first tested the dsRNA-internalization capability of the S2 and S2R+ cells from our stock. For this, we employed a commonly used Luciferase assay. Briefly, cells were transfected with plasmids expressing the Firefly and *Renilla* luciferases. Next, dsRNA targeting the Firefly luciferase sequence (dsFluc) or an unrelated control gene sequence (dsCtl: dsRNA with GFP sequence) were added to the growth medium. Firefly luciferase activity was quantified as an indirect measure of dsRNA internalization. Cells which internalize higher levels of dsRNA will exhibit less Firefly luciferase activity. *Renilla* luciferase level is used for normalization purposes. Compared to dsCtl-treated controls, we observed a decrease in Firefly luciferase activity when dsFluc was added to S2R+ cells (Supplementary Fig. 1a). These results indicate that S2R+ cells, but not S2 cells, are able to internalize dsRNA in the presence of FBS.

Considering that S2R+ cells are persistently infected with DCV, Drosophila A virus (DAV), Drosophila X virus (DXV), and Flock House virus (FHV)(32), we wondered if the infection status of these cells could account for this difference in dsRNA internalization capacity. To test if S2 cells infected with viruses can internalize dsRNA, we repeated the Luciferase assay using two different S2 cell lines previously established in our laboratory that are persistently infected with FHV (S2p FHV) or FHV and DAV (S2p FHV+DAV)(33) (Supplementary Fig. 1a). We observed a small, but significant decrease in Firefly luciferase activity in S2p FHV cells treated with dsFluc compared to those treated with dsCtl. However, no differences in Firefly luciferase activity were observed between dsFluc- and dsCtl-treated S2p FHV+DAV cells. These results suggest that, even if the infection status of the cells might have an impact on dsRNA internalization capacity, viral infection is not the primary determinant of their dsRNA internalization ability.

To avoid the confounding effects of virus infection, we decided not to use S2R+ cells but instead to develop a model for dsRNA internalization in non-infected S2 cells. We routinely culture S2 cells in Schneider cell culture medium supplemented with 10% FBS. Considering that FBS prevents dsRNA internalization by S2 cells, we adapted our uninfected S2 cell line (hereafter referred to as S2naive) to growth in Insect-XPRESS^TM^ protein-free medium, which does not contain FBS. We then tested dsRNA internalization in the newly adapted cells (hereafter referred to as S2Xpress). First, we used the Luciferase assay to compare the dsRNA internalization abilities of S2naive cells (grown in FBS-containing media) and S2Xpress cells (grown without FBS). We found a significant decrease in Firefly luciferase activity for S2Xpress cells treated with dsFluc compared to dsCtl. As we previously observed, dsFluc treatment did not significantly alter Firefly luciferase activity in S2naive cells grown in the presence of FBS (Fig. 1a). Furthermore, dsRNA internalization was dose-dependent in S2Xpress cells (Fig. 1b). We also found that substituting Insect-XPRESS^TM^ protein-free medium for 10% FBS-Schneider medium during a 4 h incubation with dsRNA in S2Xpress cells had little effect on the internalization, given that significant silencing was still observed in these conditions (Fig. 1c). This suggests the presence of a long-lasting difference between S2Xpress and S2naive cells that allows S2express cells to internalize dsRNA even in the presence of FBS. As expected, the presence of FBS had a much greater impact on dsRNA internalization for S2naive cells (Fig. 1c). We note that while some silencing was observed for S2naive cells in the absence of FBS, silencing levels were minute compared to those observed in S2Xpress cells (Fig. 1c).

**Figure 1:**
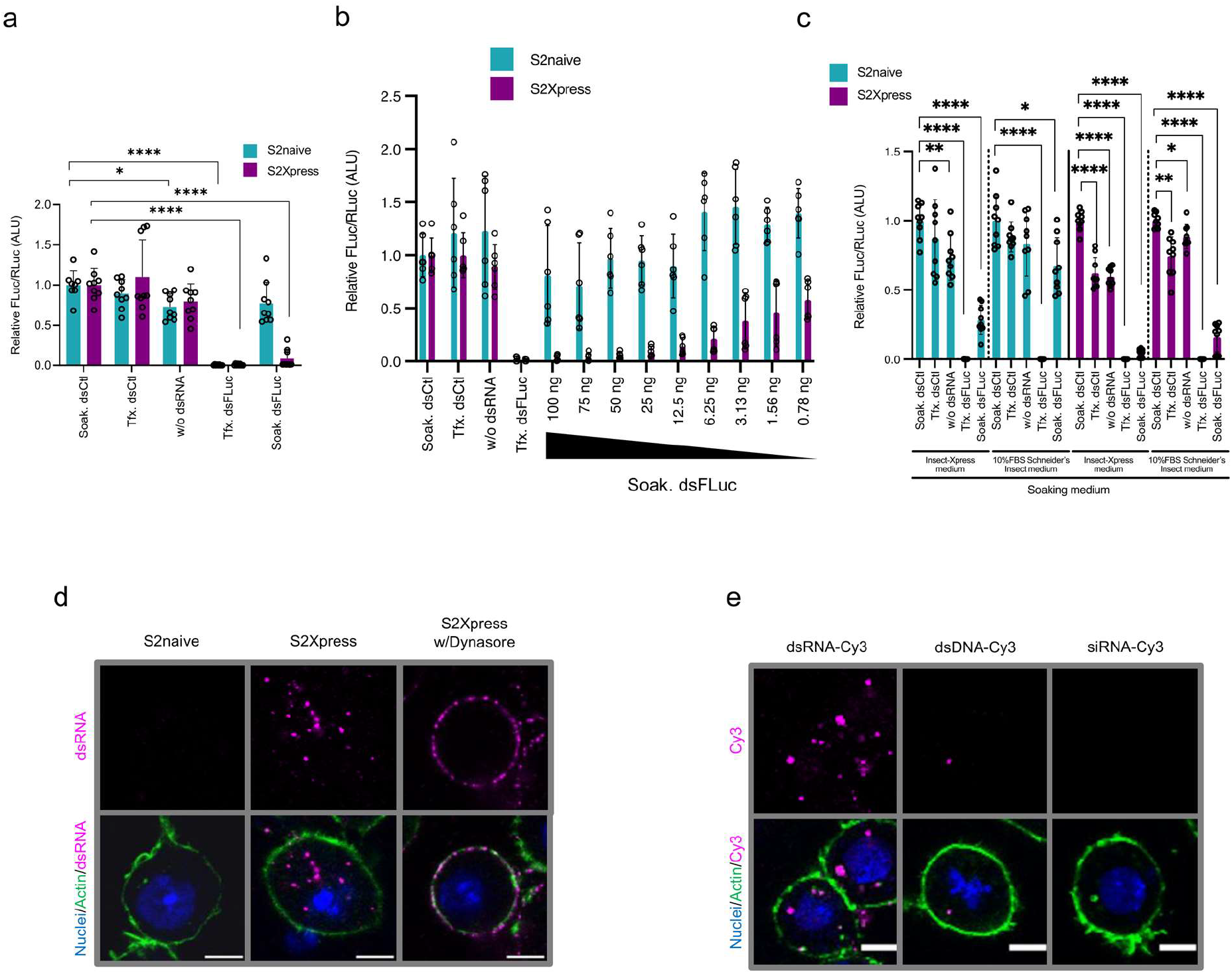
S2Xpress cells internalize dsRNA. (**a**) Silencing of Firefly luciferase by internalized dsRNA. Cells were co-transfected with plasmids expressing Firefly and *Renilla* luciferase. 24 h post-transfection, 50 ng of dsRNA targeting Firefly Luciferase (dsFLuc) or GFP (dsCtl) was added to the medium (soaking). For positive controls, dsRNAs and plasmids were co-transfected. Luciferase activity was measured 24 h after induction. Firefly luciferase values were normalized based on *Renilla* luciferase values. Data are from 3 independent experiments (n=8-9) and indicate mean ± SD of Firefly/*Renilla* ratio relative to Soak. dsCtl. Welch’s ANOVA was used to detect significant differences compared to Soak. dsCtl. (**b**) Dose-dependent silencing of Firefly luciferase. Cells were transfected as in (a) and the indicated amounts of dsRNA were added during soaking. Data are from 2 independent experiments (n=6) and indicate mean ± SD of Firefly/*Renilla* ratio relative to Soak. dsCtl. Welch’s ANOVA was used to detect significant differences compared to Soak. dsCtl. p-values are indicated in Supplementary Figure 1e. (**c**) Effects of soaking medium on dsRNA internalization. Experiments were done as in (a) using the indicated soaking medium for 4 h. Data are from 3 independent experiments (n=9) and indicate mean ± SD of Firefly/*Renilla* ratio relative to Soak. dsCtl. Welch’s ANOVA was used to detect significant differences compared to Soak. dsCtl. (**d**) Internalization of dsRNA evaluated by fluorescence confocal imaging. Cells were soaked with 30 ng of Cy3-labeled dsRNA (dsFLuc) (magenta) for 40 min. Actin and nuclei are in green and blue, respectively. Dynasore was added for 1 h before dsRNA soaking. (**e**) Specificity of nucleic acid internalization by S2Xpress cells. Cy3-labeled dsRNA (dsFLuc), dsDNA (FLuc), or siRNA (siGAPDH) (magenta) was added during soaking. Experiments were performed as in (d). ALU: arbitrary light units; Soak.: soaking; Tfx.: transfection. Confocal images were taken at 630x magnification. Scale bars represent 5 µM. p-values < 0.05 were considered significant. *p<0.05; **p<0.01; ****p<0.0001.

We next visualized internalization of dsRNA labeled with Cy3 (dsRNA-Cy3) using fluorescence confocal imaging. As before, we found that S2Xpress cells, but not S2naive cells, internalized dsRNA-Cy3 (Fig. 1d). Receptor-mediated endocytosis was previously suggested to be involved in dsRNA internalization in S2 cells(20, 21). To determine if this was also the case for S2Xpress cells, we tested the effect of Dynasore, an inhibitor of dynamin-dependent endocytosis, on the internalization of dsRNA using our dsRNA-Cy3 internalization assay. To this end, we first treated the cells with Dynasore prior to addition of dsRNA-Cy3 to the culture medium. While dsRNA-Cy3 was detected as a punctuated pattern within the cytoplasm of S2Xpress cells, dsRNA-Cy3 was localized to the cell surface when cells were treated with Dynasore (Fig. 1d). Furthermore, dsRNA internalization by S2Xpress cells was not sequence-dependent, since dsRNA internalization was still evident when we used a different dsRNA (Supplementary Fig. 1d). Finally, we found that internalization of nucleic acids by S2Xpress cells was restricted to dsRNA, since when we performed the internalization assay using dsDNA-Cy3 or siRNA-Cy3, we only saw a few spots or no spots inside the cells, respectively (Fig. 1e). Together these results show that S2Xpress cells can specifically internalize dsRNA from their environment to trigger the siRNA pathway. Since S2Xpress cells were obtained after adaptation of S2naive cells to a protein-free medium, and because only S2Xpress cells can internalize dsRNA, comparison of S2naive cells with S2Xpress cells provided us with an appropriate model system to search for proteins involved in dsRNA binding and internalization in *D. melanogaster*.

### Complementary proteomic approaches identify cell surface proteins with dsRNA binding abilities

Given that S2Xpress cells, but not S2naive cells, can internalize dsRNA, we hypothesized that differences in the composition of the plasma membranes of these two cell types might underlie differences in dsRNA internalization ability.

Two scavenger receptors, SR-CI and Eater, have been previously described to mediate the internalization of dsRNA in S2 cells(21). We tested if they affected internalization in our model. We were not able to produce dsRNA corresponding to SR-Cl since we could not obtain an RT-PCR product from S2Xpress RNA (Supplementary Fig. 2a). This precluded us from performing our dsRNA-based silencing assay for SR-Cl. For Eater, we tested dsRNA internalization by high-content fluorescence quantification and confocal imaging but we didn’t see an effect in S2xpress cells when we silenced this protein (Supplementary Fig. 2b-c). Since we could not confirm the roles of SR-CI or Eater in naked dsRNA uptake, we hypothesize that a different determinant of dsRNA internalization is present at the surface of S2Xpress cells, but not S2naive cells. To evaluate differences in cell surface components, we purified the cell surface proteins from both cell lines and determined their identity by liquid chromatography-mass spectrometry (LC-MS/MS) (Fig. 2a). Clear differences in the protein compositions of the plasma membranes between the two cell lines were observed by SDS-PAGE of the purified membrane proteins followed by silver staining (Supplementary Fig. 3a). After analyzing the cell surface protein purifications to determine protein identity, we were able to identify 37 proteins in extracts from S2naive cells and 102 proteins in extracts from S2Xpress cells, respectively (Fig. 2b, Supplementary Table 1). Only 27 of these proteins were present in both cell types. A cellular component analysis showed a high percentage of known plasma membrane proteins detected for both cell types, validating the purification protocol (Fig. 2c). For S2xpress cells, a high percentage of mitochondrion proteins was also detected. We also found some cytoplasmic and nuclear proteins in both cell types, which we speculate could be due to the presence of dead cells, or co-elution of binding partners of the plasma membrane proteins.

**Figure 2.**
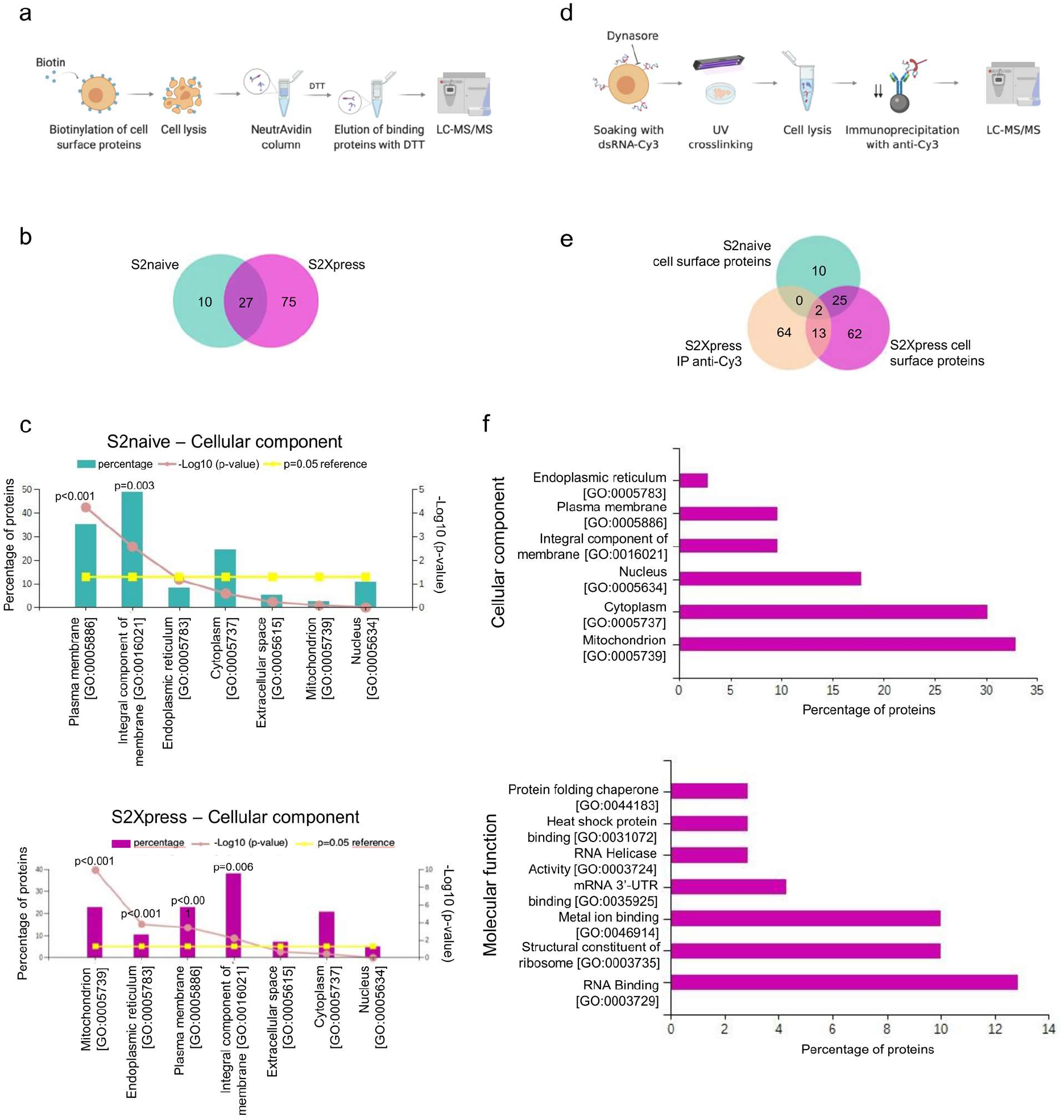
Proteomics of cell surface proteins and dsRNA binding proteins. (**a**) Scheme of the protocol used to purify cell surface proteins from S2naive and S2Xpress cells. The Pierce™ Cell Surface Biotinylation and Isolation Kit was used to biotinylate and purify cell surface proteins. Protein identification was done by LC-MS/MS. (**b**) Venn diagram showing proteins identified from S2naive and S2Xpress cells by the protocol shown in (a). (**c**) Cellular component analysis of the proteins identified by the protocol in (a) for both cells types (upper panel = S2naive, lower panel = S2Xpress). The percentages of identified proteins corresponding to the principal cell parts are shown. Statistical analysis was performed using the FunRich software(65) and p-values were calculated using a hypergeometric test. (**d**) Scheme of the protocol used to purify dsRNA-binding proteins from S2Xpress cells by immunoprecipitation (IP). Dynasore was used to prevent internalization, but not binding of Cy3-labeled dsRNA to the cell membrane. dsRNA-Cy3 was crosslinked to proteins by UV irradiation (254 nm) and IP was performed using an anti-Cy3 antibody. Precipitated proteins were identified by LC-MS/MS. (**e**) Venn diagram showing proteins identified in S2naive and S2express cells using the different proteomic protocols. (**f**) Cellular component and molecular function analysis of proteins identified by IP of dsRNA-Cy3-protein complexes from S2Xpress cells. The percentages of proteins corresponding to principal cell parts (upper panel) and relevant molecular functions (lower panel), are shown. Venn diagrams were prepared and cellular component and molecular function analyses were done with the FunRich software(65). Protocol schemes were created with BioRender.com.

To specifically purify the dsRNA-binding proteins at the cell surface we developed a state-of-the-art immunoprecipitation protocol based on the CLIP assay (CrossLinking and Immunoprecipitation) (Fig. 2d). We used Dynasore to inhibit the internalization of dsRNA-Cy3 by S2Xpress cells, thereby sequestering a high concentration of dsRNA-Cy3 at the cell surface. Next, we UV-crosslinked dsRNA-Cy3 to interacting proteins and performed an immunoprecipitation (IP) using an anti-Cy3 antibody that we found to have high specificity for dsRNA-Cy3 (Supplementary Fig. 3b). LC-MS/MS analysis of proteins purified from S2Xpress cells with the anti-Cy3 antibody allowed us to identify 79 proteins (Fig. 2e, Supplementary Table 2). When we compared the identified proteins with the ones found in the cell surface pull-down assay, we found that 13 of them were also found on the surface of S2Xpress cells, and only 2 were found on the surfaces of both cell types (S2naive and S2Xpress). Cellular component analysis showed that several of the identified proteins corresponded to mitochondrion proteins and plasma membrane proteins (Fig. 2f) as well as several nuclear proteins. This could be explained by the fact that dsRNA added to the medium seemed to bind to some dying cells present during the incubation (Supplementary Fig. 3c). In addition, co-elution of binding partners of dsRNA-binding proteins could explain the presence of cytoplasmic proteins; though it is possible that some of them are indeed undescribed plasma membrane proteins. Overall, when examining the molecular functions of the identified proteins, we found a high percentage of RNA binding proteins, indicating that the IP protocol was successful in pulling-down proteins binding to dsRNA.

Together, these two proteomic approaches allowed us to confirm that there are differences in the compositions of the cell surfaces of S2naive and S2Xpress cells, and to obtain a list of potential cell surface dsRNA-binding proteins. Of note, SR-CI and Eater were not immunoprecipitated by either of the two approaches.

### A functional screen of dsRNA-binding protein candidates identified Hsc70-4 as a protein with a role in dsRNA internalization

We next performed an *in silico* analysis of the proteins identified by our two proteomic approaches to select candidates to test in S2Xpress cells. As a selection strategy, we considered protein localization, type of protein, and presence in the results of both proteomic strategies. Through this process, we produced a list of 22 candidate proteins to test for their possible roles in mediating dsRNA internalization (Fig. 3a). To this end, we performed a high-content imaging screen based on silencing candidate proteins by transfection with candidate-gene-specific dsRNA followed by incubation with nonspecific dsRNA-Cy3. We quantified total Cy3 intensity, the number of Cy3 spots inside the cytoplasm, and the size of the spots as an indicator of dsRNA-Cy3 internalization. We successfully identified one candidate, Hsc70-4 (CG4264), that impaired the internalization of dsRNA when silenced (Fig. 3a-b, Supplementary Fig. 4a-c). We confirmed that silencing of Hsc70-4 by amplification of the full-length dsRNA target region within the candidate transcript by RT-PCR (Supplementary Fig. 4d). For Hsc70-4, for example, RT-PCR band intensity was much weaker for cells treated with dsHsc70-4 compared to cells treated with dsCtl. Furthermore, confocal imaging analysis confirmed that when Hsc70-4 was silenced, there was a decrease in the internalization of dsRNA-Cy3 (Fig. 3c). Notably, our proteomics assays identified Hsc70-4 as both a dsRNA-binding protein and a cell surface protein in S2Xpress cells, but not in S2naive cells. When we compared the expression levels of Hsc70-4 between S2naive and S2Xpress cells by RT-qPCR, we found that S2naive cells had significantly greater expression of Hsc70-4 than S2Xpress cells (Fig. 3d). Nevertheless, this result shows mRNA levels and does not necessarily reflect protein levels. Overall, these results highlight Hsc70-4 as a key factor in dsRNA internalization in S2Xpress cells and suggest that the difference between S2naive cells and S2Xpress cells regarding Hsc70-4 could relate to its localization, binding partners, or post-translational modifications.

**Figure 3.**
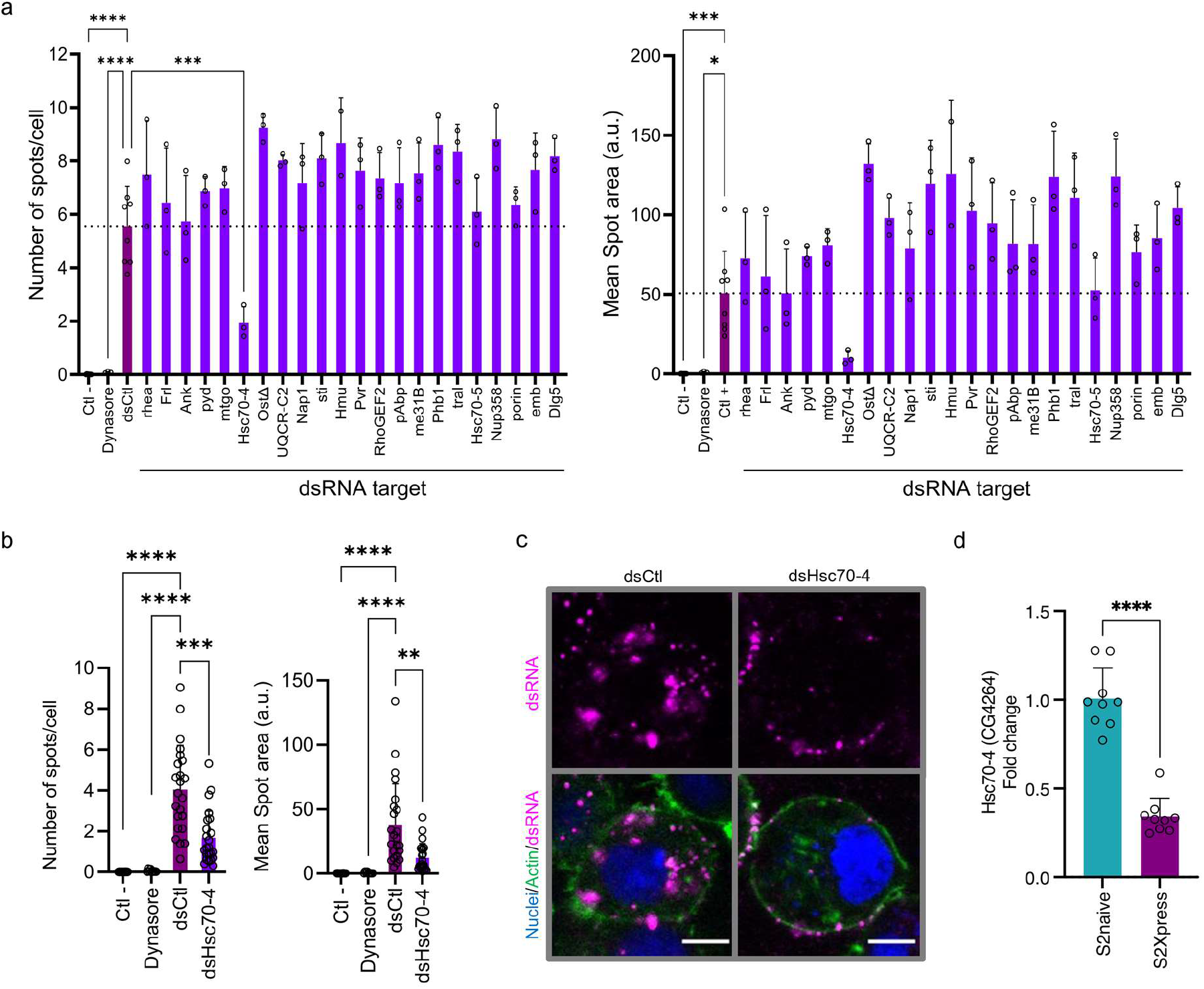
High content screen of selected candidates. (**a**) Candidate genes were silenced in S2Xpress cells by transfection with gene-specific dsRNA (dsRNA target) or nonspecific dsRNA (Ctl-, dsCtl, Dynasore) followed by soaking with Cy3-labeled nonspecific dsRNA (dsFLuc) for 30 min. For Dynasore wells, Dynasore was added for 10 min prior to soaking. Plates were imaged on an Opera Phenix High content microscope (630x magnification). Cy3 intensity, the number of Cy3 spots/cell, and the mean spot area were quantified with Columbus software. The histogram shows the mean ± SD of spots/cell (left panel) and mean spot area (right panel). A one-way ANOVA followed by Dunnett’s post-hoc tests was used to detect significant differences compared to dsCtl for both panels (Ctl- and dsCtl n=8; dsCandidates and Dynasore n=3, dsHmu n=2). p-values are for both panels are indicated in Supplementary Figure 4b. (**b**) High content analysis to determine the effects of silencing Hsc70-4 (CG4264) on dsRNA internalization. Experiments were performed as in (a). Histograms show mean ± SD of Cy3 spots/cell and mean spot area compared to dsCtl. Data are from 3 independent experiments (Ctl-n=24; Dynasore n=9; dsCtl n=23, dsHsc70-4 n=23). Welch’s ANOVAs followed by Dunnett’s T3 post-hoc test were used to detect significant differences compared to dsCtl (**c**) Confocal imaging was performed to confirm the findings shown in (b). S2xpress cells were transfected with dsHsc70-4 or dsCtl for 72 h followed by soaking with 30 ng of Cy3-labeled dsRNA (magenta) for 40 min. Actin and nuclei are shown in green and blue, respectively. Representative images from 3 experiments are shown. Confocal images were taken at 630x magnification. Scale bars represent 5 µM. (**d**) Comparison of the expression levels of Hsc70-4 between S2naive and S2Xpress cells by RT-qPCR. Rp49 was used as a housekeeping gene. Data are from 3 independent experiments and the mean ± SD of fold change relative to S2naive cells is shown. An unpaired t-test was used to detect significant differences (n=9). a.u.: arbitrary units, p-values < 0.05 were considered significant. *p<0.05; ***p<0.001; ****p<0.0001.

### Recombinant Hsc70-4 localizes to the cell surface

To better understand the role of Hsc70-4 as a mediator of dsRNA internalization, we cloned the protein fused to a V5 tag at the C-terminus in a *Drosophila* expression plasmid (hereafter referred to as pHsc70-4). The sequence and insertion site was confirmed by sequencing. This protein, referred to as recombinant Hsc70-4 (rHsc70-4), had the expected molecular size (Fig. 4a). Furthermore, when S2Xpress cells were co-transfected with pHsc70-4 and dsHsc70-4, we were not able to detect the recombinant protein, indicating that the sequence was correct and that silencing of rHsc70-4 by dsHsc70-4 was effective. Next, we transfected S2naive and S2Xpress cells with pHsc70-4 to determine the localization of rHsc70-4. We found that localization was similar in both cell types, with some cytoplasmic localization and mostly with accumulation at the cell edges (Fig. 4b).

**Figure 4.**
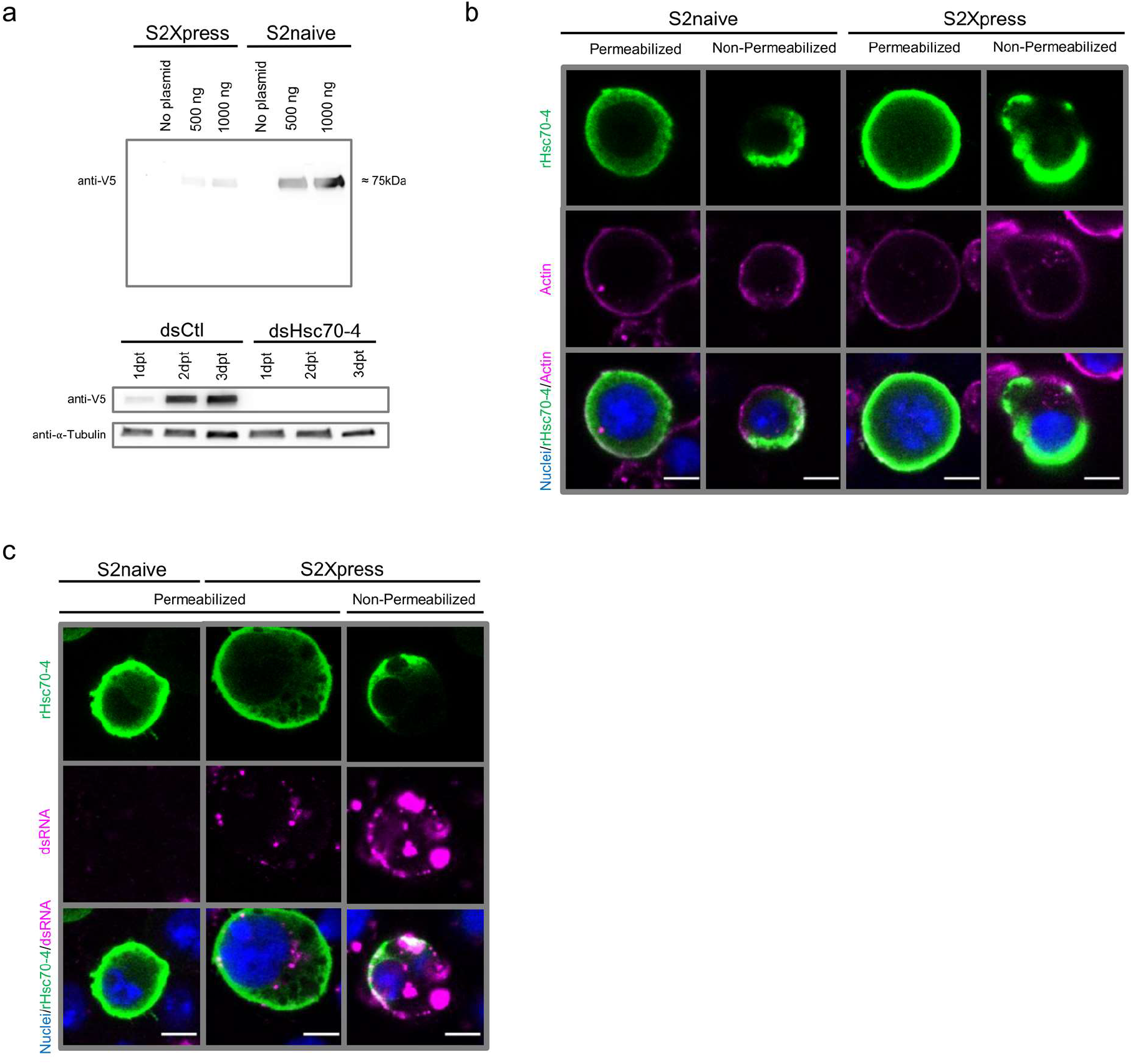
Cell surface localization of rHsc70-4. (**a**) The Hsc70-4 coding sequence was cloned into the pAc5-V5/His *D. melanogaster* expression plasmid to generate pHsc70-4. S2naive and S2Xpress cells were transfected with 0, 500, or 1000 ng of pHsc70-4 or without plasmid, and incubated for 48 h (upper panel) or 1, 2, and 3 days (lower panel). For lower panel, S2Xpress cells were co-transfected with 500 ng of pHsc70-4 and 100 ng of dsHsc70-4 or dsCtl (dsFLuc) to check silencing at the protein level. RIPA buffer was used to extract proteins. Equal volumes of samples were loaded on an SDS-PAGE gel (upper panel). For lower panel, 5 µg of proteins were loaded per lane. After electrophoresis, immunoblotting was performed with an anti-V5 antibody. α-Tubulin was used as a loading control (lower panel). The band detected by the anti-V5 antibody corresponds to the predicted molecular weight of rHsc70-4 (74.7 kDa). (**b**) S2naive and S2Xpress cells were transfected with 100 ng of pHsc70-4 for 48 h to determine cellular localization of rHsc70-4. Cells were then fixed and blocked/permeabilized with 10% normal goat serum (NGS)-0.2% Triton X-100 or 10% NGS for non-permeabilized wells. rHsc70-4 was detected by immunofluorescence with anti-V5 (green). Actin and nuclei are shown in magenta and blue, respectively. (**c**) S2naive and S2Xpress cells were transfected with 100 ng of pHsc70-4 for 48 h followed by soaking with 30 ng of Cy3-labeled dsRNA (dsFLuc) for 40 min. Cells were then fixed and blocked/permeabilized with 10% NGS-0.2% Triton X-100 or 10% NGS for non-permeabilized wells. rHsc70-4 was detected by immunofluorescence with anti-V5 (green). Nuclei are in blue. Confocal images were taken at 630x magnification. Scale bars represent 5 µM. dpt: days post transfection.

To determine if rHsc70-4 was present at the cell surface, we performed non-permeabilization assays, where the inability of antibodies to pass through the plasma membrane during the staining helps to determine if the target protein is present at the cell surface. Our results showed that rHsc70-4 was located at the cell surface, with an extracellular face in both S2naive and S2Xpress cells (Fig. 4b). Despite the cell surface localization of rHsc70-4 in in S2naive cells, expression of this protein was not sufficient to induce internalization of dsRNA (Fig. 4c). This is not surprising since we have previously found that S2naive cells do express Hsc70-4. Notably, the cell membrane localization of rHsc70-4 was preserved in mammalian cells, suggesting a conserved role for this protein at the cell surface (Supplementary Fig. 5a-b). Hsc70-4 does not present a predictable transmembrane domain(s). Its localization at the cell surface suggests that this is a peripheral membrane protein that anchors to the cell membrane by interacting with the lipid bilayer. Our results confirm that Hsc70-4 is in fact present at the cell surface and thus could be acting as a cell surface receptor or co-receptor for dsRNA.

### rHsc70-4 binds dsRNA *in vitro*

Several different domains have been identified as dsRNA binding domains in proteins, including the αβββα fold commonly known as dsRNA binding domain (dsRBD), the helicase domain, and the nucleotidyltransferase (NTase) domain(34, 35). Hsc70-4, does not present a previously identified dsRNA binding domain. Thus, we next tested if rHsc70-4 can bind dsRNA *in vitro*. rHsc70-4 was commercially produced by expression of the recombinant protein in *E. coli* followed by His-tag-based purification. Purified rHsc70-4 was incubated at different concentrations with dsRNA-Cy3 and binding of rHsc70-4 to dsRNA-Cy3 was assessed by electrophoretic mobility shift assays (EMSAs). We found that rHsc70-4 binds dsRNA *in vitro,* with increasing concentrations of rHsc70-4 resulting in decreased mobility of dsRNA-Cy3 compared to dsRNA-Cy3 alone (Fig. 5a). Moreover, the fact that the electrophoretic shift gradually increased with increasing concentrations of rHsc70-4 until reaching a full shift may indicate that more than one molecule of rHsc70-4 can bind to each molecule of dsRNA-Cy3. A competitor assay with unlabeled dsRNA was performed to confirm binding specificity. Here, we observed increased mobility for dsRNA-Cy3 in the presence of competitor dsRNA compared to the assay without competitor dsRNA, indicating that unlabeled dsRNA displaced labeled dsRNA (Fig. 5a). Furthermore, by using a dsRNA with a different sequence as a probe, we confirmed that binding of dsRNA to rHsc70-4 was not sequence-dependent (Fig. 5b). Moreover, the electrophoretic mobility of dsDNA-Cy3 or siRNA-Cy3 was not altered by incubation with rHsc70-4 (Fig. 5c-d). These results confirm that rHsc70-4 can bind dsRNA *in vitro*, and that this binding is specific for dsRNA.

**Figure 5.**
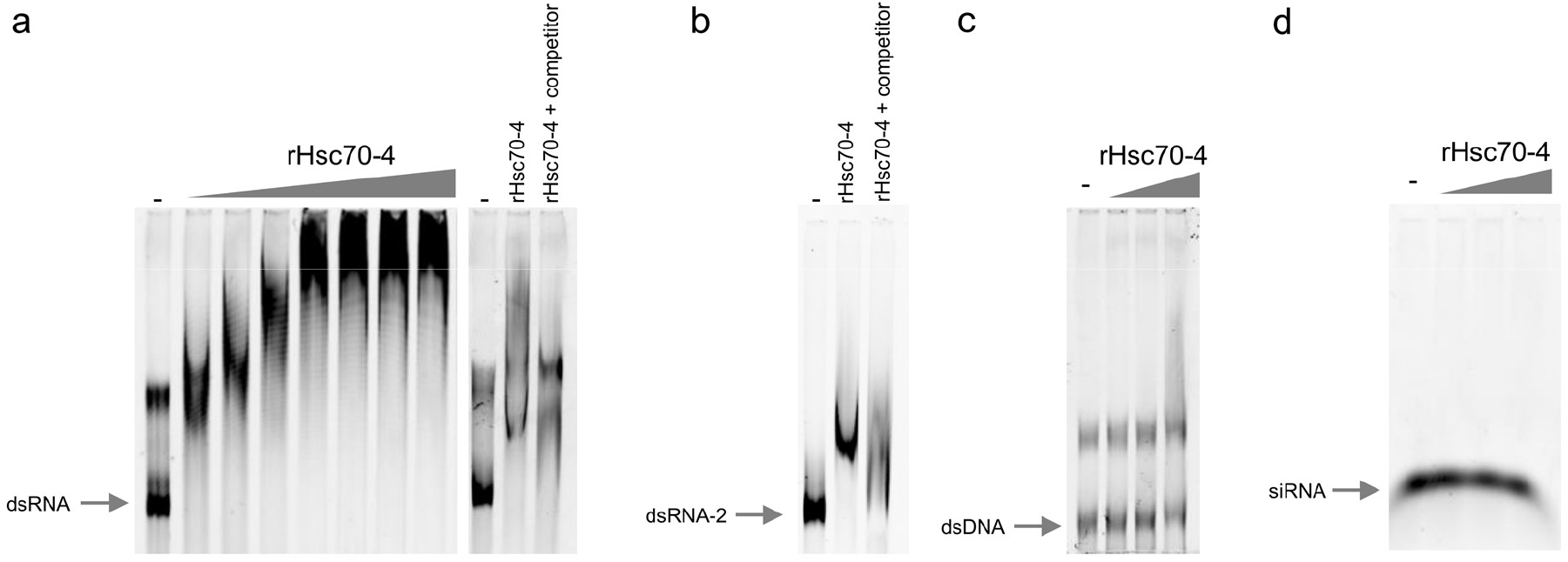
rHsc70-4 binds dsRNA, but not dsDNA or siRNA. (**a**) To test the ability of rHsc70-4 to bind dsRNA, different concentrations of rHsc70-4 (0.25 µM, 0.5 µM, 1 µM, 2µM, 3 µM, 4 µM and 5 µM) were incubated with Cy3-labeled dsRNA (dsFluc, 0.76 nM) for 30 min at 25°C and the electrophoretic shift of Cy3-labeled dsRNA was evaluated by native PAGE for 45 min on ice at 200 V in a 4% polyacrylamide gel followed by fluorescence detection using a Typhoon^TM^ FLA 9000 (left panel). dsRNA binding by rHsc70-4 was visualized by a shift in the mobility of Cy3-labeled dsRNA incubated with rHsc70-4 compared to Cy3-labeled dsRNA alone (first lane). In a competition assay (right panel), unlabeled dsRNA and Cy3-labeled dsRNA were incubated with rHsc70-4 (1 µM) in a 10:1 unlabeled:labeled dsRNA ratio. An increase in the mobility of Cy3-labeled dsRNA confirmed that unlabeled dsRNA displaced the labeled dsRNA. (**b**) To test if binding of rHsc70-4 to dsRNA was sequence dependent, a Cy3-labeled dsRNA with a different sequence was used (dsRNA-2 corresponds to dsCG6647). rHsc70-4 (2µM) was incubated with Cy3-labeled dsRNA-2 (0.76 nM) for 30 min at 25°C. The PAGE gel was run for 45 min and developed as in (a). Binding of rHsc70-4 to dsRNA-2 was visualized by a shift in the mobility of dsRNA-2 incubated with rHsc70-4 compared to dsRNA-2 alone (first lane). The competition assay was done as in (a) using dsRNA-2. (**c**) Cy3-labeled dsDNA (FLuc, 0.76 nM) was incubated with different concentrations of rHsc70-4 (0.5 µM, 1 µM, 2µM) as in (a). The PAGE gel was run for 80 min and developed as in (a). (**d**) Cy3-labeled siRNA (siGAPDH, 0.76 nM)) was incubated with different concentrations of rHsc70-4 (0.5 µM, 1 µM, 2µM) as in (a). The PAGE gel was run for 20 min and developed as in (a). All EMSA experiments were performed twice, giving similar results. The first lane in all gels corresponds to Cy3-labeled nucleic acid (dsRNA, dsDNA, or siRNA) incubated without rHsc70-4.

## Discussion

To elucidate the process of dsRNA internalization by insect cells, we sought to identify key components involved in the internalization of naked extracellular dsRNA in *D. melanogaster*. For this purpose, we utilized an *in vitro* cell model based on the adaptation of S2 cells to grow in a serum-free medium that gave us the necessary resolution and tractability to examine this very specific step of whole organism systemic immunity. Adaptation of S2 cells to growth in serum-free medium was necessary, as we and others have found that S2 cells are able to internalize dsRNA in the of FBS (supplementary Fig. 1a)(29, 36, 37). This suggests that a component of FBS may prevent dsRNA binding and/or internalization by blocking or inhibiting cell-surface proteins involved in dsRNA internalization or by sequestering dsRNA. Interestingly, we found that, while FBS does have a small impact on the ability of S2Xpress cells to internalize dsRNA, the presence FBS is not sufficient to prevent dsRNA internalization by S2Xpress cells (Fig. 1C). This suggests that S2Xpress cells underwent an intrinsic and long-lasting change compared to S2naive cells that allows them to bind and internalize dsRNA even in the presence of FBS.

The first step of dsRNA internalization is binding of dsRNA to a cell surface protein. Given their different dsRNA internalization capacities, we compared the cell surface protein compositions of S2naive and S2Xpress cells. As expected, a significant percentage of the proteins identified by our proteomic approaches were categorized as membrane proteins. Although the total number of proteins identified was less than expected, we hypothesize that the proteins we did identify may represent the more abundant and/or accessible proteins within the plasma membrane. Interestingly, even though some proteins were present at the surface of both cell types, the majority were unique to the surface of each cell type. This confirms that there are intrinsic differences in the compositions of the plasma membranes of S2naive and S2Xpress cells. Surprisingly, a significant percentage of the proteins we identified in the S2Xpress cells were mitochondrion proteins. This finding may reflect the changes undergone by these cells during adaptation to growth in protein-free media. Several reports show that mitochondrion can translocate to the plasma membrane in yeast and mammalian cells, and interact with the extracellular milieu(38, 39). Moreover, one study showed that the plasma membrane can interact with mitochondrion as a response to cellular stress by translocating caveolins to facilitate adaptation(40). Given that S2Xpress cells are grown without FBS, it is conceivable that they may experience stress due to nutrient deficiency and that changes in the composition of the plasma membrane may have allowed them to adapt to these conditions.

While our initial proteomic approach based on subcellular localization gave us valuable information about the protein composition of the surfaces of S2naive and S2Xpress cells, we wanted to add an additional layer of information to our analyses by identifying proteins present in dsRNA-protein complexes at the cell surface. For this purpose, we utilized the CLIP technique, which is widely used to study RNA-protein interactions(41–43). Our state-of-the-art CLIP-based immunoprecipitation protocol was designed to specifically isolate dsRNA-protein complexes present at the cell surface during dsRNA internalization. Indeed, a high percentage of these proteins were already identified as RNA binding proteins, including the DEAD-box RNA helicase Belle, which is involved in RNAi(21). Furthermore, Argonaute-2 (Ago-2), one of the main components of the RNAi pathway(16, 44), was also identified. Although our protocol was designed to sequester dsRNA at the cell surface by using Dynasore to inhibit internalization, inhibition of internalization was not 100% effective and some dsRNA was expected to become internalized. Thus, it is not surprising that we identified some intracellular proteins known to interact with dsRNA. In fact, our identification of intracellular RNA-binding proteins such as Ago-2 validates the effectiveness of our immunoprecipitation protocol. Together, our complementary proteomic approaches identified 15 dsRNA-binding proteins that are present at the surface of S2Xpress cells.

For our functional screening, we decided to consider all proteins as possible candidates, regardless of whether they contain a known dsRNA binding domain. Our screen of the selected candidates successfully identified one protein, heat-shock cognate 70-4 (Hsc70-4), as necessary for dsRNA internalization. Hsc70-4 is a member of the large Heat-shock protein family(45). The heat-shock cognate proteins (Hscs) differ from the more well-known heat-shock proteins (Hsps) in that they are constitutively expressed(46, 47). The Hsp70 family is comprised of a number of highly homologous proteins primarily known for their roles as chaperones, although they are involved in a wide variety of other cellular processes, including clathrin-mediated endocytosis, protein quality control, RNAi, viral attachment and entry, antiviral defense, and neurotransmitter exocytosis(48–52). They contain a highly conserved amino-terminal ATPase domain, a substrate-binding domain, and a less conserved carboxy-terminal domain(53). Because members of the Hsc70 family display high sequence similarity, it is sometimes assumed that their roles are somewhat interchangeable. Our own experiments show that this is not always the case, since another of the candidate proteins tested during our high-content screen, Hsc70-5 (CG8542; 52% amino acid sequence identity compared to Hsc70-4), was not found to be involved in dsRNA internalization.

Hsc70-4 is a highly conserved protein within the Animalia phylum. It is known to have a crucial role in the nervous system as a synaptic chaperone(51, 54). It is also involved in autophagy and protein aggregation(54, 55). Interestingly, experimental data show that Hsc70-4 is involved in clathrin-mediated endocytosis(53). Considering this, one may hypothesize that the role of Hsc70-4 during dsRNA internalization is mainly through the intracellular endocytic step. However, Hsc70-4 was previously found to participate specifically in the uncoating of clathrin-coated vesicles (CCVs) during endocytosis(53). Our microscopy-based internalization assay was designed to quantify dsRNA within the interior of the plasma membrane, thus, we would have considered dsRNA as internalized even if it was inside CCVs (but not released into the cytosol due to Hsc70-4 silencing). This suggests that Hsc70-4 may have an additional role at the cell surface during internalization of dsRNA. This is supported by our observations that rHsc70-4 accumulates at the cell surface, presents an extracellular face, and directly interacts with dsRNA. Thus, our results suggest that Hsc70-4 could act as a cell surface dsRNA receptor or co-receptor.

Our results indicate that Hsc70-4 is necessary for dsRNA internalization, but perhaps not sufficient. S2naive cells, which do not internalize dsRNA under standard growth conditions, express this protein and rHsc70-4 accumulates at the cell membrane in these cells. Nevertheless, we did not detect Hsc70-4 in our proteomics analysis of cell surface proteins in S2naive cells. It is possible that S2naive cells express Hsc70-4 at the cell surface, but at levels too low for its detection by our proteomics approach. It is also possible that another protein inhibits the detection of cell surface Hsc70-4 in S2naive cells. Importantly, our protein localization assays were done with a recombinant protein and may not necessarily reflect the expression or localization patterns of endogenous Hsc70-4. Moreover, different isoforms (7 isoforms already identified; source: FlyBase) or post-translation modifications could give different functions to Hsc70-4 in S2Xpress cells compared to S2naive cells. Finally, it is possible that Hsc70-4 may exhibit different protein-protein interactions in S2naive cells compared to S2Xpress cells in such a way that it can mediate internalization of dsRNA in S2Xpress cells, but not in S2naive cells.

How can Hsc70-4 act to internalize dsRNA? A recent report shows that the mammalian orthologue of this protein, HSPA8, acts as a co-factor receptor at the cell surface(56). Specifically, HSPA8 was found to directly interact with Porcine reproductive and respiratory syndrome virus at the cell surface and was required for virus attachment and entry(56). Interestingly, the authors showed that an additional protein, CD163, was also required for viral entry. Furthermore, HSPA8 was found to have a dual role during viral entry, as the protein directly binds to viral glycoproteins and also participates in clathrin-mediated endocytosis. Thus, these findings, together with our experimental data allow us to hypothesize a similar role for Hsc70-4 during dsRNA binding and internalization in *D. melanogaster*. Moreover, a co-factor role for Hsc70-4 in dsRNA internalization could explain why S2 cells have been shown to internalize only long dsRNA, as opposed to siRNA (21 bp)(20). One can hypothesize that if two cell surface factors, one being Hsc70-4, are necessary for dsRNA binding and internalization, shorter dsRNAs may not be long enough to bind to both proteins.

Several insects internalize extracellular dsRNA from the extracellular environment. In this context, mosquitos are of particular relevance since they serve as vectors of several human pathogenic viruses (arboviruses)(57). Interestingly, injection of dsRNA into the mosquito body cavity is sufficient to silence cognate gene expression(58, 59). In honey bees, one study showed that oral acquisition of dsRNA targeting the Israeli acute paralysis virus is sufficient to confer resistance to the treated bees(60, 61). The capacity of insects to internalize dsRNA has facilitated a new RNAi-based pest-control technology in agriculture, commonly termed “host-induced gene silencing”(60, 62, 63). Since this is a promising approach that could eventually prevent substantial agricultural and economic losses, it is vital to acquire a thorough knowledge of the mechanism of action of the internalization of dsRNA, including its target tissues, cell internalization and processing, and transmission. The present work is a starting point that will facilitate the study of dsRNA internalization mechanisms through the identification of one of the key elements of this process. To our knowledge, this is the first report showing that Hsc70-4 directly binds dsRNA and that it is present at the cell membrane with an extracellular face. Further studies are needed to characterize the dsRNA binding domain of Hsc70-4, its binding partners, and its structure. Moving forward, the study of endogenous Hsc70-4 as well as its role *in vivo* will provide more detailed information on the specific mechanisms by which Hsc70-4 promotes dsRNA internalization.

## Methods

### Cells, plasmids and antibodies

*D. melanogaster* Schneider 2 (S2naive) cells (Invitrogen), S2R+(30) (*Drosophila* Genomics Resource Center), S2p FHV and S2p FHV+DAV(33) were cultured at 25°C in Schneider’s Insect Medium (GIBCO), supplemented with 10% heat-inactivated fetal bovine serum (GIBCO), 2 mM L-glutamine (GIBCO), 100 U/ml penicillin (GIBCO) and 100 mg/ml streptomycin (GIBCO). S2Xpress cells were cultured in Insect-XPRESS^TM^ protein-free medium (LONZA, Belgium) supplemented with 100 U/ml penicillin and 100 mg/ml streptomycin (GIBCO). S2naive and S2Xpress cells were checked by next-generation sequencing of small RNAs for infection with CrPV, DAV, DCV, DXV, FHV, Nora virus, Sigma virus, American nodavirus, and Drosophila birnavirus. HeLa cells (ATCC) were cultured at 37°C with 5% CO_2_ in DMEM + GlutaMAX medium (GIBCO) supplemented with 10% heat-inactivated fetal bovine serum, 2 mM L-glutamine, 100 U/ml penicillin and 100 mg/ml streptomycin. pMT/V5-HisB (Invitrogen) expressing either Firefly or *Renilla* Luciferase under the control of a copper-inducible promoter were previously generated (20). The coding region for Hsc70-4 (CG4264) was cloned into pAc5.1/V5-His A (Invitrogen), and into pcDNA™6/myc-His A (Invitrogen), from an amplicon produced by RT-PCR from S2Xpress RNA. Assembly was done with NEBuilder HiFi DNA Assembly Master Mix (E2621L, NEW ENGLAND BioLabs). Correct insertion of the amplicon was confirmed by sequencing. The following antibodies were used: anti-Cy3/Cy5 (ab52060, Abcam), anti-dsRNA J2 (10010500, SCICONS), anti-V5 Tag (46-0705, Invitrogen), anti-6X His Tag (ab18184, Abcam), anti-α-tubulin (T5168, Sigma-Aldrich), anti-Mouse IgG (HRP) (ab6728, Abcam), anti-Mouse IgG (Alexa Fluor 488) (A11029, Invitrogen), anti-Mouse IgG (Alexa Fluor 555) (A21422, Invitrogen).

### Adaptation of S2 cells to Insect-Xpress medium

To generate the S2Xpress cells, S2naive cells were gradually adapted to the Insect-XPRESS protein-free medium. Briefly, S2naive cells were grown until confluency in a T25 flask in normal growth medium (Schneider’s Insect Medium, supplemented with 10% FBS), and then transferred into a T75 flask, adding 5 ml of Insect-Xpress medium. 4 days after, 5 ml of Insect-XPRESS medium were added. At day 8, cells with medium were transferred into a T150 flask, and 15 ml of Insect-XPRESS medium were added. At day 11, 15 ml of cell suspension were transferred into a new T150 flask, and 15 ml of Insect-XPRESS medium were added. From this point forward, cells were adapted to the Insect-XPRESS medium, and were passaged only using this medium.

### dsRNA production

dsRNA was produced by *in vitro* transcription using the MEGAscript T7 Transcription kit (AM1334, Invitrogen) following the manufacturer’s guidelines. For dsRNAs targeting endogenous genes, cDNA produced with SuperScript™ II Reverse Transcriptase (100004925, Invitrogen) from S2Xpress RNA was used as PCR template to amplify the target regions using primers flanked by the T7 promoter. For dsFLuc (Firefly Luciferase, GL3) and dsGFP, plasmids were used as templates (pMT-GL3 and pAc5.1-GFP). dsRNA concentration was quantified on a Qubit 3 fluorometer. All dsRNAs produced against candidates were 450-600 bp in length. dsGL3: 557 bp; dsGFP: 714 bp, dsCG6647: 508 bp. The full list of primers can be found in Supplementary Table 3.

### Nucleic acid labeling with Cy3

The *Silencer* siRNA Labeling kit – Cy3 (AM1632, Invitrogen) was used to label dsRNA (dsFLuc, dsGFP, dsCG6647), DNA (FLuc) or siRNA (GAPDH, from the kit) with Cy3 following the kit’s protocol. Labeling was confirmed by an electrophoretic shift of the purified product compared to unlabeled probe on a 1.5% agarose gel and by immunofluorescence with the anti-dsRNA antibody, J2 (Supplementary Fig. 1b-c).

### Fluorescence microscopy

S2naive and S2Xpress cells were seeded in an 8-chamber Nunc LabTek II that had previously been coated with Poly-L-Lysine (P4832, Sigma-Aldrich). Cells were then incubated for 24 h. For silencing experiments, cells were transfected with dsRNA using the Effectene Transfection reagent (301427, QIAGEN) and incubated for 72 h. For endocytosis inhibition experiments, Dynasore (D7693, Sigma-Aldrich) was added to a final concentration of 100 μM for 10-60 min at 25°C. dsRNA, DNA or siRNA labeled with Cy3 (to a final concentration of 0.76 nM) was added and cells were incubated at 25°C for 30-40 minutes. Cells were washed with PBS, fixed in 4% paraformaldehyde for 20 min, and blocked/permeabilized in 2% BSA-0.2% Triton X-100. Actin was visualized with Oregon Green™ 488 Phalloidin (O7466, Invitrogen). For immunofluorescence with J2 and anti-Cy3 antibodies, blocking/permeabilization was done with 10% normal goat serum-0.2% Triton X-100 followed by an overnight incubation at 4°C with a primary antibody (J2: 1:500; anti-Cy3: 1:500). Slides were incubated with a secondary antibody (anti-Mouse IgG-Alexa Fluor 488: 1:1000) for 1 h, and no actin staining was performed. For immunofluorescence to detect rHsc70-4, cells were transfected with pAc5.1-rHsc70-4-V5/His (S2 cells) or with pcDNA6-rHsc70-4-myc/His (Hela cells) and incubated for 48 h before fixing. For Hela cells, transfection was performed with Lipofectamine 3000 (L3000015, Invitrogen). Staining was done as previously described, except that Triton X-100 was omitted in non-permeabilized wells. Primary antibodies were used at 1:1000 dilution for anti-V5 Tag, and 1:400 for anti-6X His. Actin was visualized with Alexa Fluor™ 555 Phalloidin (A34055, Invitrogen) (except when dsRNA-Cy3 was used). Nuclei were counterstained with DAPI. Vectashield H-1000 (Vector Laboratories, Burlingame, CA) was used as mounting medium. Imaging was done on a Leica TCS SP5 confocal microscope at 630x magnification (S2 cells) or 400x magnification (Hela cells), and images were processed in Fiji(64).

### Luciferase assays

Cells were seeded on 96-well culture plates and transfected with plasmids pMT-GL3 (Firefly Luciferase, 12 ng/well) and pMT-*Renilla* (*Renilla* Luciferase, 3 ng/well), and with dsRNA (10 ng/well) when indicated, using the Effectene Transfection reagent (301427, QIAGEN). After 24 h, the medium was changed, and dsFLuc or dsCtl (dsGFP) (50 ng/well, or the indicated amounts) was added (soaking) to allow internalization and posterior silencing of FLuc. After 24 h, plasmid expression was induced by adding 10 mM CuSO_4_. The next day, cells were lysed, and measurement of Firefly and *Renilla* Luciferase activity was performed using the Dual-Luciferase Reporter Assay System (E1960, Promega) in a GLOMAX microplate luminometer. Firefly Luciferase values were normalized to *Renilla* Luciferase values. For analysis of the effect of the medium on internalization, the medium was changed to the indicated medium prior to soaking and then soaking was performed for 4 h. Following soaking, the medium was changed back to the appropriate growth medium for each cell type. The experiments for Supplementary Figure 1 were performed with minor modifications: after transfection, cells were incubated 48 h before soaking with dsRNA. After plasmid induction with CuSO_4_, cells were incubated another 48 h before cells lysis.

### Purification and identification of cell surface proteins

S2naive and S2Xpress cells were seeded in T75 flasks and incubated until the next day, when they were collected and counted. 2.4 x10^7^ cells were diluted in 15 ml of growth medium. Biotinylation and purification of biotinylated proteins were done following the Pierce™ Cell Surface Biotinylation and Isolation Kit’s (A44390, Thermo Scientific) protocol. Samples without biotin were used as negative controls to identify nonspecific protein precipitation. Samples were eluted in 200 μl of elution buffer. 50 μl of purified proteins were used for proteomic analysis and a separate 50 μl aliquot was used for gel analysis. 6x Laemmli SDS sample buffer (J60660, Alfa Aesar) (with β-mercaptoethanol) was added to the samples and the samples with Laemmli buffer were boiled for 5 min at 100°C. For proteomic analysis, samples were loaded on a 7.5% SDS-PAGE gel (BioRad) before in-gel proteolysis. Briefly, gel bands corresponding to proteins were excised and washed and then the proteins were reduced and alkylated (DTT 10 mM final, 2h, 37°C iodoacetamide, 50 mM final, 30 min in the dark at room temperature). After dehydration, protein proteolysis was done with 50 ng of LysC-trypsin (Promega) at 37°C, overnight. The resulting proteolytic peptides were extracted from the gel by incubating twice for 15 min in 100 µL 1% aq. TFA and sonication, followed by one incubation of 15 min at 37°C in 50 µL acetonitrile. Peptides were desalted using C18-filled tips (Ziptip^TM^ C18, Millipore), eluted in 10 µL of 0,1% TFA in 50% acetonitrile, dried, and dissolved in 8 µL buffer A. The digest (6 µL of peptides) was injected on a capillary reversed-phase column (C18 Acclaim PepMap100 A, 75 µm i.d., 50-cm length; Thermo Fisher Scientific) at a flow rate of 220 nL/min, with a gradient of 2% to 40% Buffer B in Buffer A in 60 min (Buffer A: H_2_O/acetonitrile(ACN)/ FA 98:2:0.1 (v/v/v); Buffer B: H_2_O/ACN/FA10:90:0.1 (v/v/v)). The MS analysis was performed on a Qexactive HF mass spectrometer (Thermo Fisher Scientific) with a top 10 data dependent acquisition method: MS resolution 70,000, mass range 400-2,000 Da, followed by MS/MS on the ten most intense peaks at resolution 17,500, with a dynamic exclusion for 10 s. Raw data was processed using Proteome Discoverer 2.4 (Thermo Fisher Scientific). The database search was done with the Mascot search engine (Matrix Science Mascot 2.2.04) on a *D. melanogaster* protein databank (20986 entries). The SwissProt databank 2020_05 (563,552 entries) was used to assess contamination with human proteins. The following parameters were used: MS tolerance 10 ppm; MS/MS tolerance 0.02 Da; tryptic peptides; up to two miscleavages allowed; partial modifications: carbamidomethylation C, oxidation (M), deamidation (NQ). Proteins identified by at least two high confidence peptides (FDR<0.1%) were validated. Further analysis of the identified proteins was done on FunRich software(65). For SDS-PAGE, samples were loaded on a 4-15% polyacrylamide gradient gel. After electrophoresis, the gel was fixed with 40% ethanol-10% acetic acid-50% H_2_O and silver staining was performed as previously described(66).

### Immunoprecipitation of dsRNA binding proteins

S2Xpress cells were seeded on 100 mm culture plates (1 x 10^7^ cells/10 ml/plate) and incubated for 24 h at 25°C. Then the medium was changed and the endocytosis inhibitor Dynasore (D7693, Sigma-Aldrich) was added (final concentration 100 μM) and cells were incubated for 1 h at 25°C. Next, dsRNA (dsFLuc) labeled with Cy3 was added (10 μg/plate) and soaking was performed for 45 min at 25°C. For negative control plates, unlabeled dsRNA was used. Following soaking, plates were washed in cold PBS and UV-crosslinked (254 nm) in 3 ml of PBS (without Mg/Ca) at 300 mJ/cm^2^ on ice in a Stratalinker UV 1800 crosslinker. Cells were scraped from the plates, pelleted by centrifugation, and lysed in SDS lysis buffer (0.5% SDS, 50 mM Tris, 1 mM EDTA, 1 mM DTT, pH 8, 1x Protease inhibitor (11 873 580 001, Roche)) at 65°C for 5 min. Next, RIPA correction buffer was added (62.5 mM Tris, 1.25% NP-40 substitute, 0.625% sodium deoxycholate, 2.25 mM EDTA, 187.5 mM NaCl, pH 8, 1x Protease inhibitor), and samples were passed through QIAshredder columns (79654, QIAGEN) twice, following the manufacturer’s recommendations. Following this, the samples were incubated with 4 μg of antibody (anti-Cy3) overnight at 4°C on a rotator. The next day, the Dynabeads™ Protein A Immunoprecipitation Kit (10006D, Invitrogen) was used to pull-down dsRNA-protein complexes binding to the antibody, following the kit’s protocol. Briefly, samples were incubated with the magnetic beads for 1 h at RT. Next, samples were diluted and washed 5 times with RIPA buffer (50 mM Tris, 1% NP-40 substitute, 0.5% sodium deoxycholate, 0.1% SDS, 2 mM EDTA, 150 mM NaCl, pH 8, 1x Protease inhibitor (911 873 580 001, Roche)) prior to elution. To elute proteins, the samples were first treated with RNase III for 1 h at 37°C, and supernatants were collected. Next, the beads were treated with 6x Laemmli SDS sample buffer (J60660, Alfa Aesar) (with 10% β-mercaptoethanol), boiled for 5 min at 100°C and the supernatants were collected. Proteomics analysis was done as detailed in the previous section. Samples collected after RNase III treatment had only a few proteins, indicating that the elution was not effective. Thus, only samples collected after boiling in sample buffer were considered. Duplicates were performed for each treatment. Negative control samples were used to discard proteins that were nonspecifically precipitated. Analysis of the identified proteins was done using the FunRich software(65).

### High-content imaging screen

Cells were seeded on black CellCarrier Ultra microplates (6055302, PerkinElmer) and transfected with 10 ng/well of dsRNA targeting the candidates using the Effectene Transfection reagent (301427, QIAGEN). dsCtl and Dynasore (D7693, Sigma-Aldrich) wells were transfected with dsCtl (dsFLuc). After 72 h of incubation, the medium was changed and Dynasore was added to corresponding wells (final concentration 100 μM) for 10 min at 25°C. Next, 30 ng/well of dsRNA (dsFLuc) labeled with Cy3 was added. Cells were then incubated for 30 min at 25°C, fixed in 4% paraformaldehyde for 20 min, and blocked/permeabilized in 2% BSA-0.2% Triton X-100 for 30 min. The cytoplasm was visualized with DiO dye (from Vybrant™ Multicolor Cell-Labeling Kit; V22889, Molecular Probes) and nuclei were counterstained with DAPI. Imaging was performed on an Opera Phenix microscope (PerkinElmer) at 630x magnification. The Columbus software was used to design a script and analyze the images. The script was designed to quantify the median intensity of Cy3 and the number of spots of Cy3 in the cytoplasm per well. Further analysis was done using GraphPad Prism 9. For the median fluorescence intensity, negative control wells were used to subtract baseline intensity values.

### RT-qPCR

Cells were seeded in 24-well plates and incubated for 24 h. RNA extraction was performed with TRIzol™ Reagent (15596026, Ambion), and RNA concentration was quantified using a NanoDrop One (Thermo Scientific). Equal amounts of RNA were treated with RQ1 DNase (M610A, Promega) and were used to produce cDNA with Maxima H Minus First Strand cDNA Synthesis Kit (K1682, Thermo Scientific) using random primers. Quantitative PCR was done with Luminaris Color HiGreen qPCR Master Mix, low ROX (K0374, Thermo Scientific) on a QuantStudio™ 7 Flex Real-Time PCR System (Applied Biosystems). Rp49 was used as a housekeeping gene. ΔCt values were obtained by subtracting the Ct value for Rp49 from the Ct value of the corresponding gene and ΔΔCt values were obtained by further subtracting the geometric mean ΔCt value for the control condition (S2naive). Results are shown as fold change relative to S2naive. Primer sequences can be found in Supplementary Table 3.

### RT-PCR to confirm silencing

To confirm silencing of Hsc70-4 (CG4264) by dsHsc70-4, cells were transfected with dsHsc70-4 with the Effectene Transfection reagent (301427, QIAGEN) and incubated for 72 h. For the control condition, cells were transfected with dsCtl (dsFLuc). RNA was extracted and cDNA was produced as previously described, with the modification that Oligo(dT)_18_ primers were used. PCR was performed with DreamTaq DNA Polymerase (EP0702, Thermo Scientific) using primers flanking the dsRNA targeting region. PCR products were loaded on a 1% agarose gel with ethidium bromide. After electrophoresis, the gel was developed on a Gel Doc XR+ Imaging System (Bio-Rad). Rp49 was used as a loading control. Primers sequences can be found in Supplementary Table 3.

### Western blot

Cells were seeded on 6-well plates and transfected with 500 ng or 1000 ng of pAc5.1-rHsc70-4-V5/His (S2 cells) or with 100 ng of pcDNA6-rHsc70-4-myc/His (Hela cells). After 48 h, cells were washed with cold PBS and proteins were extracted with RIPA buffer ((50 mM Tris, 1% NP-40 substitute, 0.5% sodium deoxycholate, 0.1%SDS, 2 mM EDTA, 150 mM NaCl, pH 8, 1x Protease inhibitor (11 873 580 001, Roche)). Samples were incubated on ice for 20 min, and centrifuged for 20 min at 4°C 13000 *g*.

Supernatants were collected. To confirm expression of the recombinant protein, equal volumes of samples were boiled in XT Sample Buffer (161-0791, Bio-Rad) with β-mercaptoethanol for 5 min, and loaded on a 4-20% Mini-PROTEAN® TGX Stain-Free™ Protein Gel (4568094, Bio-Rad). PageRuler™ Prestained Protein Ladder (26616, Thermo Scientific) was used as a molecular weight ladder. After electrophoresis, the gel was activated and then transferred for 30 min to a Trans-Blot Turbo Mini 0.2 µm nitrocellulose membrane (1704158, Bio-Rad) on a Trans-Blot Turbo transfer system (BioRad). Total proteins were visualized by UV on a Gel Doc XR+ Imaging System (Bio-Rad). Membranes were blocked with 5% non-fat dry milk in PBS-0.05% Tween-20 and incubated overnight with primary antibody at 4°C (anti-V5 Tag: 1:5000, anti-6X His: 1:5000, anti-α-tubulin: 1:5000). Washes were done with PBS-0.05% Tween. Membranes were incubated for 1.5 h with secondary antibody (anti-Mouse IgG (HRP): 1:5000), developed with SuperSignal™ West Pico PLUS Chemiluminescent Substrate (34580, Thermo Scientific), and imaged on a G:BOX Chemi XL (Syngene). For silencing experiments, S2Xpress cells were co-transfected with 500 ng of plasmid and 100 ng of dsHsc70-4 or dsCtl (dsFLuc) and incubated for 1, 2, or 3 days. The protein concentration in the samples was quantified with the Pierce™ BCA Protein Assay Kit (23227, Thermo Scientific) and equal amount of proteins were loaded on the gel. α-Tubulin was used as a loading control.

### Electrophoretic mobility shift assay (EMSA)

For EMSA experiments, dsRNA (dsFLuc, dsCG6647), dsDNA (FLuc), or siRNA (GAPDH) labeled with Cy3 were used as probes. The recombinant Hsc70-4 protein was produced by GenScript by expression of the recombinant protein with a 6X His tag in *E. coli*. The protein was purified with Ni resin. Labeled-probe (0.76 nM) was incubated with different concentrations of the recombinant protein in binding buffer (25 mM Tris, 50 mM NaCl. 0.1 mg/ml BSA, 31.25 mM DTT, 0.625 mg/ml tRNA *E. coli*; 20 μl reaction) for 30 min at 25°C. Native loading buffer was added after incubation (100 mM Tris, 10% glycerol, 0.0025% bromophenol blue, pH: 8). For competition assays, 10 times more of the same unlabeled-dsRNA was added to the reaction mix. EMSA gels (native 4% polyacrylamide, 0.5X TBE) were pre-run at 200 V on ice prior to loading samples in 0.5X TBE buffer. Samples were loaded and ran at 200 V on ice. Gels were imaged by fluorescence detection on a Typhoon^TM^ FLA 9000 (GE Healthcare).

### Statistical analysis

Data are represented as mean ± SD. Statistical analyses were done in GraphPad Prism version 9 (GraphPad Software, CA, USA). Two-tailed unpaired t-tests (2 groups) or one-way ANOVAs (3 or more groups) followed by Dunnett’s post-hoc tests were used to detect significant differences between groups. p-values < 0.05 were considered significant. Normality and homoscedasticity were assumed for the data. Whenever these criteria were not met, non-parametric Welch’s ANOVA tests (3 or more groups) followed by Dunnett’s T3 post-hoc tests were used. For cellular component analysis of proteomic results, A hypergeometric test was done using the FunRich software(65).

### Data availability

All data generated in this study, including proteomic results, are provided in the Supplementary Information. The mass spectrometry proteomics data have been deposited to the ProteomeXchange Consortium via the PRIDE [1] partner repository with the dataset identifier PXD034508 and 10.6019/PXD034508.

## Acknowledgements

We thank all members of the Saleh lab for insightful discussion, and Jared Nigg for critical reading and editing of the manuscript. We would like to specially thank the UtechS Photonic BioImaging (Imagopole), C2RT, Institut Pasteur, which is supported by the French National Research Agency (France BioImaging; ANR-10–INBS–04; Investments for the Future) and the Région Île-de-France (program DIM1Health), for assistance with imaging, especially Julien Fernandes, Nathalie Aulner and Anne Danckaert. We thank Felix Rey (Institut Pasteur) and Daved Fremont (Washington University) for advice on isolating membrane proteins. We also thank Lluis Quintana-Murci, Marco Vignuzzi and James Di Santo for buying the reagents needed for experiments. This work was supported by a Pasteur-Roux-Cantarini fellowship from Institut Pasteur to S.J.F., and by the European Research Council (FP7/2013-2019 ERC CoG 615220) and the French Government’s Investissement d’Avenir program, Laboratoire d’Excellence Integrative Biology of Emerging Infectious Diseases (grant ANR-10-LABX-62-IBEID) to M.-C.S. Mass spectrometry equipment was subsidized by Conseil Régional d’Île-de-France (Sesame N°10022268).

## Authors contributions

S.J.F. and M-C.S. conceived the project. S.J.F., L. T-P. and V. M. performed experimental design, development, and analysis of results. L.F. and H.B. did the transcriptomic analysis. Y.V and J.V. carried out the LC-MS/MS procedures. M-C. S. supervised the project and obtained the funding. All authors contributed to the preparation of the manuscript.

