## Supplementary Figures for "Hsc70-4 mediates internalization of environmental dsRNA at the surface of *Drosophila* S2 cells"

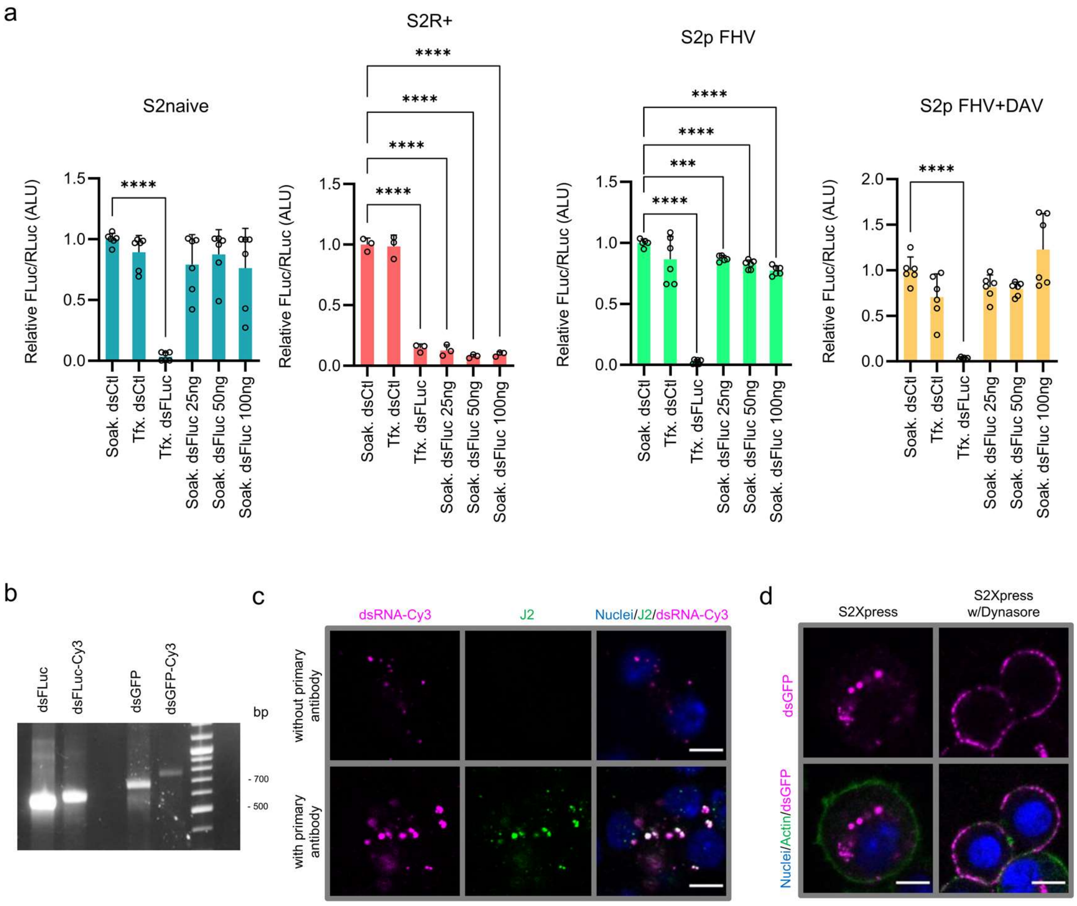


e

| Treatment | p-value (vs. Soak. dsCtl) S2naive | p-value (vs. Soak. dsCtl) S2Xpress |
|---|---|---|
| Tfx. dsCtl | ns | ns |
| w/o dsRNA | ns | ns |
| Tfx. dsFLuc | 0.0006 | 0.0002 |
| Soak. dsFLuc 100 ng | ns | 0.0003 |
| Soak. dsFLuc 75 ng | ns | <0.0001 |
| Soak. dsFLuc 50 ng | ns | 0.0003 |
| Soak. dsFLuc 25 ng | ns | 0.0001 |
| Soak. dsFLuc 12.5 ng | ns | <0.0001 |
| Soak. dsFLuc 6.25 ng | ns | <0.0001 |
| Soak. dsFLuc 3.13 ng | ns | 0.0117 |
| Soak. dsFLuc 1.56 ng | ns | 0.0493 |
| Soak. dsFLuc 0.78 ng | ns | 0.0181 |

*Supplementary Figure 1: dsRNA internalization by different S2 cell lines*. (**a**) Luciferase assays to test internalization of dsRNA by S2naive, S2R+, S2p FHV and S2p FHV+DAV cells. Cells were co-transfected with plasmids expressing Firefly and *Renilla* luciferase. After 48 h, 25, 50 or 100 ng of dsRNA targeting Firefly Luciferase (dsFLuc) or GFP (dsCtl) was added to the medium (soaking) to allow internalization and silencing of Firefly luciferase. For control experiments, dsRNAs were co-transfected with plasmids. Luciferase activity was measured 48 h after plasmid induction of Luciferase expression. Firefly luciferase values were normalized based on *Renilla* luciferase values. Data are shown as mean $\pm$ SD of Firefly/*Renilla* ratio relative to the control condition (Soak. dsCtl). Data shown are from 2 experiments for S2naive, S2p FHV, and S2p FHV+DAV (n=5-6), and form 1 experiment for S2R+ (n=3). Welch's ANOVA tests were used to detect significant differences compared to the control condition (Soak. dsCtl) for the S2naive, S2p FHV, and S2p FHV+DAV experiments. A one-way ANOVA was used for the S2R+ experiment. p-values $< 0.05$ were considered significant. (**b**) Agarose gel electrophoresis of Cy3-labeled and unlabeled dsRNA (dsFLuc and dsGFP). Image contrast was adjusted for improved visualization (**c**) Immunofluorescence of S2Xpress cells with the anti-dsRNA antibody, J2 (green) to confirm that labeling of dsRNA (dsGL3) with Cy3 (magenta) was efficient and that the signal detected by immunofluorescence with J2 was in fact dsRNA rather than free Cy3. Cells were soaked with 30 ng of dsRNA-Cy3 for 40 min. DAPI was used to stain nuclei (blue). Confocal images showed complete co-localization between Cy3 signal and J2 signal, confirming that the Cy3 signal comes from the dsRNA-Cy3. (**d**) Cy3-labeled dsRNA corresponding to the sequence of GFP (dsGFP, magenta) was used to confirm that uptake of dsRNA by S2Xpress cells was not sequence-dependent. Soaking with dsRNA was done as in (c). Actin is in green and nuclei are in blue. For Dynasore experiments, Dynasore was added for 20 min before soaking with dsRNA. (**e**) Table displaying the p-values of panel (b) from Figure 1. ALU: arbitrary light units; Soak.: soaking; Tfx.: transfection. Confocal images were taken at 630x magnification. Scale bars represent 5 μM. ***p<0.001; ****p<0.0001.

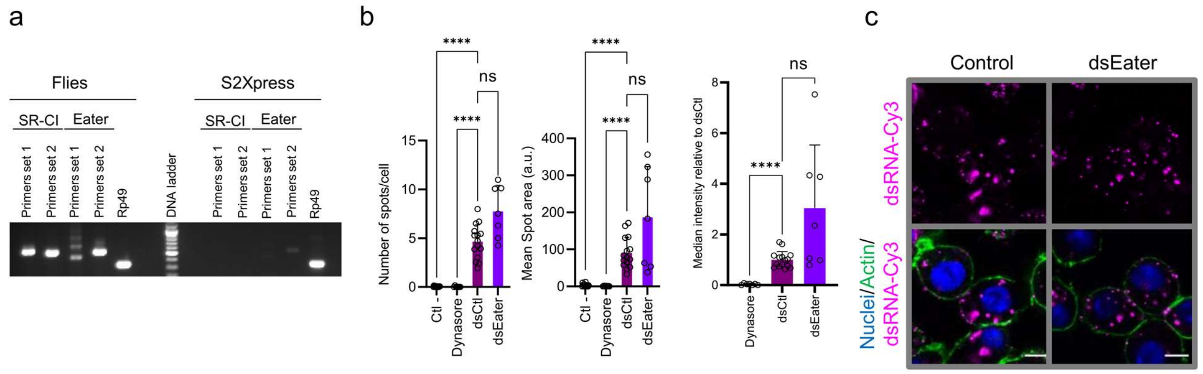


*Supplementary Figure 2.* The *SR-CI and Eater receptors are not involved in uptake of dsRNA in S2Xpress cells*. (**a**) Agarose gel electrophoresis of RT-PCRs product used as templates for *in vitro* transcription of dsRNA targeting SR-CI and Eater. Two sets of primers were used for each receptor. cDNA from $w^{1118}$ flies was used a positive control for the primers. Rp49 was used as loading and PCR control. We were not able to amplify SR-CI when cDNA from S2Xpress cells was used for PCR. The Eater PCR amplicon was purified and Eater-specific dsRNA was produced by *in vitro* transcription (dsEater). (**b**) High content imaging was used to test the effect of silencing Eater on the uptake of dsRNA. S2Xpress cells were transfected with 10 ng of dsEater to silence the receptor for 72 h. dsCtl and Dynasore wells were transfected with nonspecific dsRNA (dsFLuc). For Dynasore wells, Dynasore was added for 10 min prior to soaking with 30 ng of nonspecific Cy3-labeled dsRNA (dsFLuc) for 30 min. Plates were imaged on an Opera Phenix High content microscope at 630x magnification. A script was designed on the Columbus software to quantify Cy3 intensity and number of Cy3 spots inside the cells. Histograms show mean ± SD of the number of Cy3 spots/cell, mean spot area, and median Cy3 intensity compared to the control condition (dsCtl). Data are from 2 independent experiments pooled together (Ctl- n=15; Dynasore n=7; dsCtl n=15, dsEater n=7). For median intensity, Ctl- values were subtracted from the other conditions. Values are shown relative to dsCtl. Welch's ANOVA tests were used to detect significant differences compared to the control condition (dsCtl). p-values < 0.05 were considered significant. ****p<0.0001. (**c**) Confocal imaging was performed to test the effect of silencing Eater on the internalization of dsRNA by S2Xpress cells. S2xpress cells were transfected with dsEater or not transfected (Control) for 24 h prior to soaking with 30 ng of Cy3-labeled dsRNA (dsFLuc, magenta) for 40 min. Actin is in green and nuclei are in blue. Confocal images were taken at 630x magnification. Scale bars represent 5 μM. a u.: arbitrary units.

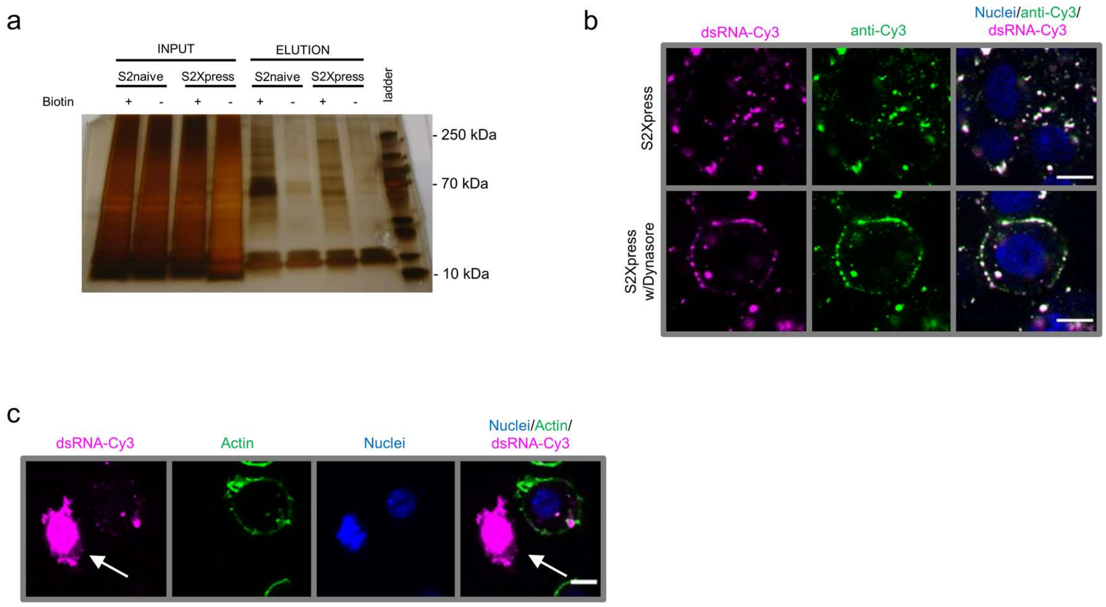


*Supplementary Figure 3. Comparison of cell surface proteins and controls for IP of dsRNA-protein complexes.* (**a**) Aliquots of proteins purified by the biotinylation of cell surface proteins of S2naive and S2Xpress (see Figure 2a) were separated on an SDS-PAGE gel and stained by silver staining. INPUT samples refer to lysed samples prior to isolation of biotinylated proteins. ELUTION samples refer to isolated proteins that were further analyzed by LC-MS/MS. Differences between S2naive and S2Xpress cells were evidenced by the different banding patterns of the ELUTION lanes. Negative control samples were prepared by not adding biotin during labeling of cell surface proteins. (**b**) Immunofluorescence with an anti-Cy3 antibody was performed to check specificity of antibody to be used for the IP of dsRNA binding proteins from S2Xpress cells. 30 ng of Cy3-labeled dsRNA (dsFLuc, magenta) was added to the cells for 40 min. Inhibition of endocytosis with Dynasore was performed by adding Dynasore for 1 h prior to soaking. Immunofluorescence was performed with an anti-Cy3 antibody (green). Nuclei are in blue. Confocal images were taken at 630x magnification. Complete co-localization of dsRNA-Cy3 and anti-Cy3, even when Dynasore was added, confirmed that this antibody is highly specific and thus a good option for IP. (**c**) Visualization of the internalization of dsRNA-Cy3 (magenta) was performed as in (b) with minor modifications. Actin was stained with phalloidin (green) and DAPI was used for nuclei (blue). The arrow indicates a dying/dead cell with high Cy3 signal. dsRNA seems to bind to these death nuclei/cells, resulting in the immunoprecipitation of nuclear proteins. Confocal images were taken at 630x magnification. Scale bars represent 5 μM.

a

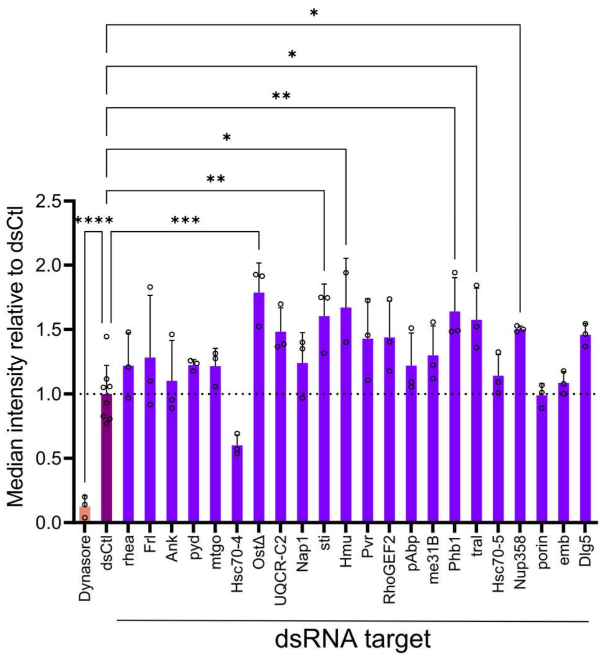


b

| Treatment | Number of spots/cell p-value vs. dsCtl | Mean Spot area p-value vs. dsCtl |
|---|---|---|
| Ctl - | <0.0001 | 0.0009 |
| Dynasore | <0.0001 | 0.0408 |
| rhea | 0.2393 | 0.9451 |
| Frl | 0.9891 | 0.9992 |
| Ank | 0.9997 | >0.9999 |
| pyd | 0.8137 | 0.9147 |
| mtgo | 0.7128 | 0.6142 |
| Hsc70-4 | 0.0004 | 0.1854 |
| OstΔ | 0.0003 | <0.0001 |
| UQCR-C2 | 0.0481 | 0.0632 |
| Nap1 | 0.5229 | 0.7109 |
| sti | 0.0363 | 0.0009 |
| Hmu | 0.0228 | 0.0023 |
| Pvr | 0.1639 | 0.0288 |
| RhoGEF2 | 0.3467 | 0.1077 |
| pAbp | 0.5278 | 0.5679 |
| me31B | 0.2166 | 0.5730 |
| Phb1 | 0.0050 | 0.0003 |
| tral | 0.0134 | 0.0056 |
| Hsc70-5 | 0.9992 | 0.9999 |
| Nup358 | 0.0021 | 0.0003 |
| porin | 0.9949 | 0.8312 |
| emb | 0.1499 | 0.3955 |
| Dlg5 | 0.0251 | 0.0206 |

c

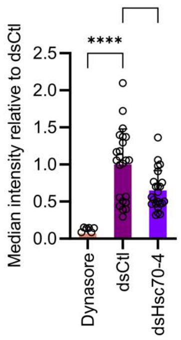


d

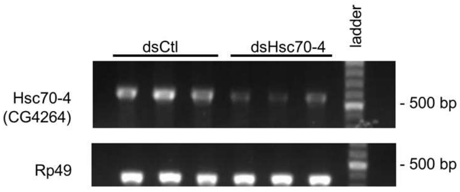


*Supplementary Figure 4. High content screen intensity and silencing confirmation*. (**a**) 22 selected candidates were silenced in S2Xpress cells by transfecting with 10 ng of candidate gene-specific dsRNA (dsRNA target) for 72 h. dsCtl and Dynasore wells were transfected with nonspecific dsRNA (dsFLuc). For Dynasore wells, Dynasore was added for 10 min prior to soaking with 30 ng of nonspecific Cy3-labeled dsRNA (dsFLuc) for 30 min. Plates were imaged on an Opera Phenix High content microscope at 630x magnification. A script was designed on the Columbus software to quantify Cy3 intensity and the number of Cy3 spots inside the cells. Histograms show mean $\pm$ SD of median Cy3 intensity relative to the control condition (dsCtl). Ctl- values were subtracted from other the conditions, and values are shown relative to dsCtl. A one-way ANOVA was used to detect significant differences compared to the control condition (dsCtl) (dsCtl n=8; dsCandidates and Dyansore n=3, dsHmu n=2). (**b**) Table shows specific p-values of panel (a) from Figure 3. (**c**) High content analysis to determine the effects of silencing Hsc70-4 (CG4264) on dsRNA internalization. The experiment was performed as in (a). Histograms show mean $\pm$ SD of median Cy3 intensity relative to the control condition (dsCtl). Ctl- values were subtracted from the other conditions and values are shown relative to dsCtl. Data are from 3 independent experiments (Dynasore n=9; dsCtl n=23, dsHsc70-4 n=23). Welch's ANOVAs followed by Dunnett's T3 post-hoc tests were used to detect significant differences compared to dsCtl (**d**) To confirm silencing of Hsc70-4 by dsHsc70-4, S2Xpress cells were transfected with 50 ng of dsHsc70-4 or dsCtl (dsFLuc) for 72 h (triplicates per condition). Next, RNA was extracted, quantified, and cDNA was produced from equal amounts of RNA using Oligo(dT)$_{18}$ primers. PCR was performed with primers flanking the dsRNA targeting region. PCR products were separated on a 1% agarose gel with ethidium bromide. Rp49 was used as a loading and PCR control. Silencing was confirmed by the decreased intensity of the bands from cells treated with dsHsc70-4. The experiment was performed twice, with similar results each time. p-values < 0.05 were considered significant. *p<0.05; **p<0.01; ***p<0.01; ****p<0.0001.

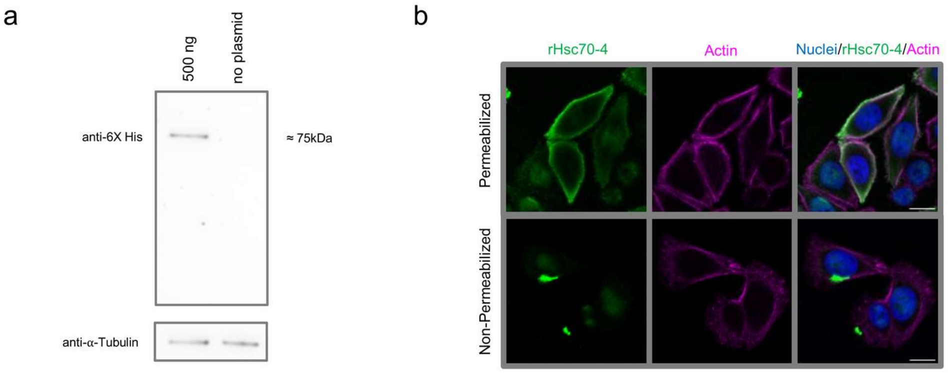


*Supplementary Figure 5. rHsc70-4 localizes to the cell surface of HeLa cells*. (**a**) The Hsc70-4 coding region was cloned into the pcDNA6-myc/His mammalian expression vector. HeLa cells were transfected with 1000 ng of this plasmid (or non-transfected for negative control) and incubated for 48 h. RIPA buffer was used to extract proteins. 4 µg of proteins were separated by SDS-PAGE followed by immunoblotting with an anti-6X His antibody. α-Tubulin was used as a loading control. The band detected with the anti-6x His antibody corresponds to the predicted molecular weight of rHsc70-4 (75.4 kDa). (**b**) HeLa cells were transfected with 100 ng of the plasmid encoding rHsc70-4 for 48 h followed by fixation and blocking/permeabilization with 10% NGS-0.2% Triton X-100 or 10% NGS for non-permeabilized wells. rHsc70-4 was detected by immunofluorescence with an anti-6X His antibody (green) overnight at 4°C. Actin is in magenta and nuclei are in blue. Confocal images were taken at 400x magnification. Scale bars represent 15 µM.
